# Relative value perception in an insect: positive and negative incentive contrasts in ants

**DOI:** 10.1101/330241

**Authors:** Stephanie Wendt, Kim S. Strunk, Juergen Heinze, Andreas Roider, Tomer J. Czaczkes

**Affiliations:** Animal Comparative Economics laboratory, Institute of Zoology & Evolutionary Biology, University of Regensburg, 93053 Regensburg, Germany; School of Business, Economics and Information Systems, Chair of Management, People and Information, University of Passau, 94032 Passau; Institute of Zoology & Evolutionary Biology, University of Regensburg, 93053 Regensburg, Germany; Department of Economics, University of Regensburg, 93053 Regensburg, Germany

**Keywords:** Incentive contrasts, successive contrasts, relative value perception, foraging, recruitment, private information

## Abstract

Humans tend to value things not on their absolute values, but relative to reference points such as former experience or expectations. People rate the quality of a new salary relative to their previous salary and the salaries of their peers, instead of appreciating its absolute value. Here, we demonstrate a similar effect in an insect: ants, which had previously experienced a low quality food source, showed higher acceptance of medium quality food (e.g. 0.1M then 0.5M; positive contrast) than if they had received the medium food all along (e.g. 0.5M then 0.5M; control), and vice versa for high expectations. Further experiments demonstrate that these contrast effects arise from cognitive rather than mere sensory or pre-cognitive perceptual causes. Pheromone deposition also correlates with perceived reward value, and ants showed successive contrasts in their pheromone deposition. Relative value perception can therefore be expected to have strong effects not only on individual behaviour, but also on collective decision-making. Contrast effects were also social: the quality of food received from other ants affected the perceived value of food found later. Value judgement is a key element in decision making, and thus relative value perception will strongly influence how animals interact with their environment.

## Introduction

We all compare options when making both large and small decisions, ranging from career choice to the choice of an evening’s entertainment. Understanding how options are compared has thus been central to the study of behaviour and economics. Theories explaining the mechanisms by which options are compared and decisions are made have a long tradition (Vlaev et al. 2011), with Expected Utility Theory (EUT) being the most widely used theory in economic models (Mankiw 2011; von Neumann and Morgenstern 1944). EUT suggests that decisions are made by evaluating and comparing the expected pay-off from each option. A rational decision maker then chooses the option resulting in the best end state (i.e. the option providing the greatest utility) (von Neumann and Morgenstern 1944).

However, over the past decades economic research on how humans make decisions has started to shift away from a view of (absolute) utility maximization towards more nuanced notions of relative utility, such as reference-dependent evaluations. Kahneman and Tversky (1979) made a major contribution to this shift by introducing Prospect Theory, suggesting that decision making is not based on absolute outcomes, but rather on relative perceptions of gain and losses. In contrast to EUT, the utility attributed to options being evaluated is determined relative to a reference point, such as the status quo or former experience (Kahneman and Tversky 1979; Parducci 1984; Tversky and Kahneman 1992; Ungemach, Stewart, and Reimers 2011; Vlaev et al. 2011). Various examples of relative value perception have been described. For example, satisfaction gained from income is perceived not absolutely, but relative to the income of others in the social reference group – such as one’s colleagues (Boyce, Brown, and Moore 2010). Overall, Prospect Theory has enriched our understanding of human decision making by conceptualizing it as more nuanced than previously assumed (Tversky and Kahneman 1974, 1981).

A similar relativistic pattern can be found in sensory judgements: Humans rated drinks containing the same sucrose concentration sweeter when they were presented with a range of lower concentrations and less sweet when higher concentrations were presented more frequently (McBride 1982; Riskey, Parducci, and Beauchamp 1979). However, these findings also match well with predictions from psychophysics, in which the link between a given stimulus strength and it’s sensation is studied (Zwislocki 2009). A key psychophysical finding is that identical stimuli are perceived as more or less intense depending on the strength of reference stimuli.

The concept of malleable value perception is not just relevant to humans. Value judgments in animals are also influenced by factors apparently independent of the absolute value of options. For example, capuchin monkeys refuse otherwise acceptable pay (cucumber) in exchanges with a human experimenter if they had witnessed a conspecific obtain a more attractive reward (grape) for equal effort (Brosnan and de Waal 2003). Rats, starlings, and ants, like humans, place greater value on things they work harder for (Aw, Vasconcelos, and Kacelnik 2011; Czaczkes, Brandstetter, et al. 2018; Lydall, Gilmour, and Dwyer 2010), and fish and locusts demonstrate state-dependent learning, wherein they show a preference for options experienced when they were in a poor condition (Aw et al. 2009; Pompilio, Kacelnik, and Behmer 2006). Roces and Núñez aimed to show that in leaf cutting ants perceived value can be influenced by other ants. Ants recruited to higher quality food sources ran faster, deposited more pheromone, but cut smaller leaf fragments, even if the food source the recruits find is replaced by a standardised food source (Roces 1993; Roces and Núñez 1993). However, in these experiments the absolute value and nature of the reference remains unclear, and indeed pheromone presence may have caused the observed behaviours without influencing the ants’ expectations or value perception at all. Critically missing from this body of work is a systematic description of value judgment relative to a reference point.

A common way in which value is judged is by either comparing two options to each other or by comparing one option to an option experienced in the past. Thus, the perceived value of an option is likely to depend strongly on the strength of contrast between both options and on whether the new option results in a relative gain or a loss. Such value-distortion by comparison effects have been studied for decades using the successive contrasts paradigm. In such experiments, animals are trained to a quality or quantity of reward which is then suddenly increased (positive incentive contrast) or decreased (negative incentive contrast) (Bentosela et al. 2009; Bitterman 1976; Couvillon and Bitterman 1984; Crespi 1942; Flaherty 1982, 1999; Mustaca, Bentosela, and Papini 2000; Weinstein 1970b). The reaction of animals towards the post-shift reward is then compared to the reaction of animals which always received the first reward and therefore did not experience a shift. Many mammals, including apes, monkeys, rats and dogs (Bentosela et al. 2009; Brosnan and de Waal 2003; Crespi 1942; Flaherty 1999; Mustaca, Bentosela, and Papini 2000; Pellegrini and Mustaca 2000; Weinstein 1970a) have been shown to respond to successive negative contrast by disrupting their behaviour compared to control animals which had not experienced a change in reward. The animals display behaviour akin to disappointment – slower running speeds to a reward (Bower 1961), depressed licking behaviour (Flaherty, Becker, and Pohorecky 1985; Vogel, Mikulka, and Spear 1968), or reward rejection (Tinklepaugh 1928).

However, unlike negative contrast effects, responses to positive successive contrast have rarely been found, even when searched for (Black 1968; Capaldi and Lynch 1967; Bower 1961; Dunham 1968; Papini et al. 2001). This may be due to three possible factors, which have the opposite effect of positive contrast and may counterbalance it: ceiling effects, neophobia, and generalization decrement (Annicchiarico et al. 2016; Flaherty 1999). Ceiling effects may occur when the performance of animals receiving a large reward is at or near a physical limit. The absence of positive contrast may then not be generated by behavioural principles, but through an artefact of experimental design (Bower 1961; Campbell et al. 1970). Neophobia may manifest itself through the reluctance to eat novel food – even if the food is of higher quality than normal (Flaherty 1999). Generalization decrement may occur when animals are trained under one set of stimuli and then tested under another. The strength of the tested response may decrease with increasing differences between the training and testing stimuli (Kimble 1961), which may then result in weaker positive contrast effects following a reward shift. Thus, the reward change itself may lead to a decrease in responding just as would any other change in context, such as a change in the brightness of the runway (Capaldi 1978; Premack and Hillix 1962).

Even though positive contrast effects proved to be hard to demonstrate in laboratory experiments, there are good theoretical reasons for expecting both positive and negative contrast effects to evolve (McNamara, Fawcett, and Houston 2013): if conditions become rich in the environment of an animal which was initially exposed to poor conditions, it should work harder than if conditions have been rich all along. This is because conditions are likely to worsen in the future and the animal should therefore use good conditions to the fullest while available. By contrast, if the animal was accustomed to rich conditions which then suddenly worsen, it should work less hard than if conditions have always been poor. In this case, rich conditions are likely to return and the animal would do better by waiting for the good conditions to return before continuing to exploit the environment. Lastly, contrast effects should be strongest in animals adapted to rapidly changing conditions, because it enhances the differential allocation of effort between favourable and unfavourable periods (McNamara, Fawcett, and Houston 2013).

Contrast effects could potentially arise without differential valuation of options; other mechanisms could also in principle produce these results: contrast effects in sensory tasks could derive from simple psychophysical mechanisms (Fechner 1860; Zwislocki 2009), and thus arise from sensory perceptual mechanisms rather than higher level cognitive processing of value. Sensory judgements are also usually made relative to reference points and through constant comparisons with former stimuli (Helson 1964; Vlaev et al. 2011). The position of the reference point in the range of stimuli may thus bias how the stimulus, and thus the value, of a post-shift reward is perceived (Zwislocki 2009). For example, the sweetness of a sucrose solution may be perceived much stronger when the reference point to which it is compared is low. Sensory satiation may also result in apparent contrast effects: the more sweetness receptors are blocked by a sweet reference solution, the fewer receptors will fire when confronted with a post-shift reward, thus making solutions taste less sweet (Bitterman 1976). A final potential driver of apparent contrast effects is related to the theoretical benefits of such behaviour described above: animals may rationally expect the pre-shift reward to be available in the future again and therefore rationally show lower acceptance towards the post-shift reward, because they are waiting for the pre-shift reward to reoccur.

The finding of contrast effects in the honey bee, until now the only invertebrate for which such behaviour was conclusively shown, led to a fourth explanation for contrast effects (Couvillon and Bitterman 1984; Bitterman 1976; Núñez 1966). Bitterman (1976) found that honey bees which were trained to a 40% sucrose solution show many feeding interruptions when experiencing a downshift to 20% sucrose. By contrast, bees which were fed on 20% throughout the whole experiment filled their crops immediately. Bees which were shifted from 20% to 40% showed no interruptions at the post-shift solution either. Apart from explaining these results as negative contrast effects, Bitterman suggested two alternative hypotheses: sensory saturation (see above) and changes in satiation level. Individuals may not only store sucrose solutions in their crop, but may also ingest small amounts of sucrose, leading to an increase of haemolymph-sugar levels. Higher blood-sugar levels negatively affect sweetness perception in humans (Mayer-Gross and Walker 1946; Melanson et al. 1999), and a similar effect could cause a post-shift solution to taste less sweet to animals trained on high sucrose concentrations. However, using an odour training paradigm, Couvillon and Bitterman (1984) found negative contrast effects in honeybees and could rule out the above alternative causes.

In this study, we investigate positive and negative contrast effects using the successive contrasts paradigm, and define the first relative value curve in an invertebrate; the ant *Lasius niger*. We then demonstrate that relative value perception arises from non-rational cognitive effects, rather than rational decision-making, physiological effects, or psychophysical phenomena. Finally, we demonstrate that information flowing into the nest can influence value perception in outgoing foragers.

## Methods and Results

### Study animals

Eight stock colonies of the black garden ant *Lasius niger* were collected on the University of Regensburg campus. The colonies were kept in 30×30×10cm foraging boxes with a layer of plaster covering the bottom. Each box contained a circular plaster nest box (14 cm diameter, 2 cm height). The colonies were queenless with around 1000-2000 workers and small amounts of brood. Queenless colonies still forage and lay pheromone trails, and are frequently used in foraging experiments (Devigne and Detrain 2002; Dussutour et al. 2004). The colonies were fed with *ad libitum* 0.5M sucrose solution and received *Drosophila* fruit flies once a week. Water was available *ad libitum.*

One sub-colony of 500 individuals was formed from each stock colony, and these eight fixed-size sub-colonies were used for our experiments. Sub-colonies were maintained identically to the stock colonies, but did not receive any *Drosophila* fruit flies to prevent brood production, and were starved four days prior to the experiments in order to achieve a uniform and high motivation for foraging (Mailleux, Detrain, and Deneubourg 2006; Josens and Roces 2000). During starvation, water was available *ad libitum*. Any ants which died or were removed from the sub-colonies were replaced with ants from the original stock colonies.

### General setup

The general setup used for all of our three experiments was identical and consisted of a 20 × 1 cm long paper-covered runway which was connected to the sub-colony’s nest box via a 40 cm long drawbridge (figure 1A). A 5mm diameter drop of sucrose solution (Sigma-Aldrich) was placed on an acetate feeder at the end of the runway (60cm from the nest). The molarity of the sucrose droplet depended on the experiment, treatment and on the ants’ number of visit to the food source.

**Fig. 1:**
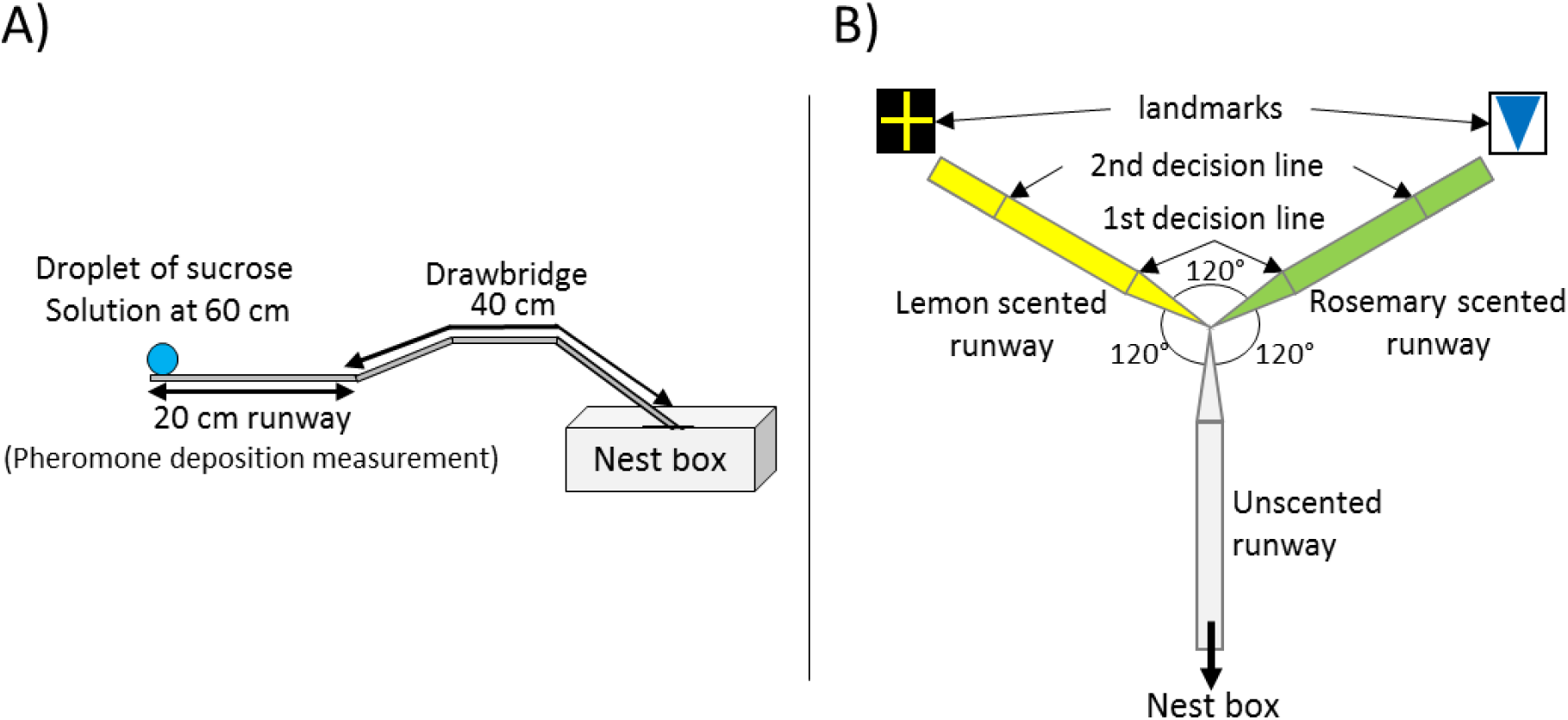
**A)** General setup used for all presented experiments. The 20 cm long runway is connected to the nest box via a 40 cm long drawbridge. The droplet of sucrose solution is placed at the end of the runway (60 cm distance to the nest). **B)** Y-maze used on the 10^th^ visit of experiment 2. All arms were 10 cm long. The arm connected to the nest box was covered with unscented paper overlays while the other two arms were covered with lemon and rosemary scented paper overlays (one scent on each side). Visual cues (landmarks) were placed directly behind the two scented arms. The first decision line was located 2.5cm from the Y-maze centre and marked the initial decision of an ant while the second decision line was located 7.5cm from the centre and marked the final decision.

To begin an experiment, the sub-colony was connected to the runway via the drawbridge. 2-4 ants were allowed onto the runway, and the first ant to reach the feeder was marked with a dot of acrylic paint on its gaster. The marked ant was allowed to drink to repletion at the food source, while all other ants were returned to the nest. As the ant drank at the droplet it was given one of three food acceptance scores. Full acceptance (1) was scored when the ant remained in contact with the drop from the moment of contact and did not interrupt drinking within 3 seconds of initial contact. Partial acceptance (0.5) was scored if feeding was interrupted within 3 seconds after the first contact with the food source, but the ant still filled its crop within 10 minutes (as can be seen by the distention of the abdominal tergites). Lastly, rejection (0) was scored if the ant refused to feed at the sucrose solution and either returned to the nest immediately or failed to fill its crop within 10 minutes.

When the ant had filled its crop or decided not to feed at the sucrose droplet, it was allowed to return to the nest. Inside the nest, the ant unloaded its crop to its nestmates and was then allowed back onto the runway for another visit. The drawbridge was now used to selectively allow only the marked ant onto the runway.

In addition to measuring food acceptance, we also measured pheromone deposition. Individual pheromone deposition behaviour correlates with the (perceived) quality of a food source (Beckers, Deneubourg, and Goss 1993; Hangartner 1970; Czaczkes, Grüter, and Ratnieks 2015). Individual ants can adapt the strength of a pheromone trail by either depositing pheromone or not, or varying the intensity of a pheromone trail through number of pheromone depositions (Hangartner 1970; Beckers, Deneubourg, and Goss 1993). Pheromone deposition behaviour in *L. niger* is highly stereotypic. To deposit pheromone, an ant briefly interrupts running to bend its gaster and press the tip of the gaster onto the ground (Beckers, Deneubourg, and Goss 1992). This allows the strength of a pheromone trail to be quantified by counting the number of pheromone depositions over the 20 cm runway leading to the feeder. Pheromone depositions were measured each time the ant moved from the food source back to the nest (inward trip), and each time the ant moved from the nest towards the food source (outward trip). Because *L. niger* foragers almost never lay pheromone when they are not aware of a food source (Beckers, Deneubourg, and Goss 1992), we did not measure pheromone depositions for the very first outward trip (visit 1). The presence of trail pheromone on a path depresses further pheromone deposition (Czaczkes et al. 2013). Thus, each time an ant had passed the 20 cm runway, the paper overlay covering the runway was replaced by a fresh one every time the ant left the runway to feed at the feeder or returned to the nest.

All experimental runs were recorded with a Panasonic DMC-FZ1000 camera to allow for later video analysis.

After each experimental run the ant was permanently removed.

Details of our statistical analysis methods and samples sizes are provided in online supplement S1.

## Experiment 1 – Defining a relative value perception curve

The aim of this of experiment was to test whether *Lasius niger* foragers value a given absolute sucrose solution concentration relative to a reference point or based on its absolute value. We used a range of 12 molarities as reference points in order to describe a value curve. To exclude effects of the researcher’s expectations on the data, the data for this experiment were collected blind to treatment (Holman et al. 2015).

### Experiment 1 - Methods

In the first two visits to the apparatus - termed the training visits - the ants’ reference point was set by allowing it to feed from a feeder at the end of the runway. The quality of the sucrose solution was varied between ants, with each ant receiving the same quality twice successively. 12 different molarities were used: 0.1, 0.2, 0.3, 0.4, 0.5, 0.6, 0.7, 0.8, 0.9, 1, 1.5 or 2M. *Lasius niger* workers learn the quality of a feeder within 2 visits (Wendt and Czaczkes 2017). On the third visit (test visit), the food source was replaced by a 0.5M sucrose solution droplet for all ants. Thus, ants trained to qualities <0.5M experienced a positive successive contrast, ants trained to >0.5M experienced negative successive contrast, and the ants trained to 0.5M constituted the control (no contrast). Food acceptance and pheromone depositions were noted for each visit, as described above.

### Experiment 1 - Results

Ants seemed to value sucrose solution droplets relative to a reference point (figure 2, table S1). While in the training visits acceptance scores increased significantly with increasing molarity of the reference quality (CLMM: 1^st^ visit: estimate= 1.18, z=6.99, p>0.001; 2^nd^ visit: estimate= 1.56, z=9.28, p<0.001, fig. 2), in the test (contrast) visit acceptance scores decreased significantly with increasing molarity of the reference quality (CLMM: estimate=-2.59, z= -13.75, p<0.001, fig. 2). Ants which were trained to very low molarities (0.1M: p<0.001) showed significantly higher acceptance of 0.5M sucrose than control ants, while ants trained to high molarities (1.5M: p<0.001, 2M: p<0.001) showed lower acceptance of 0.5M than the control group (see supplement Table S2.1 for all pairwise comparisons).

**Fig. 2:**
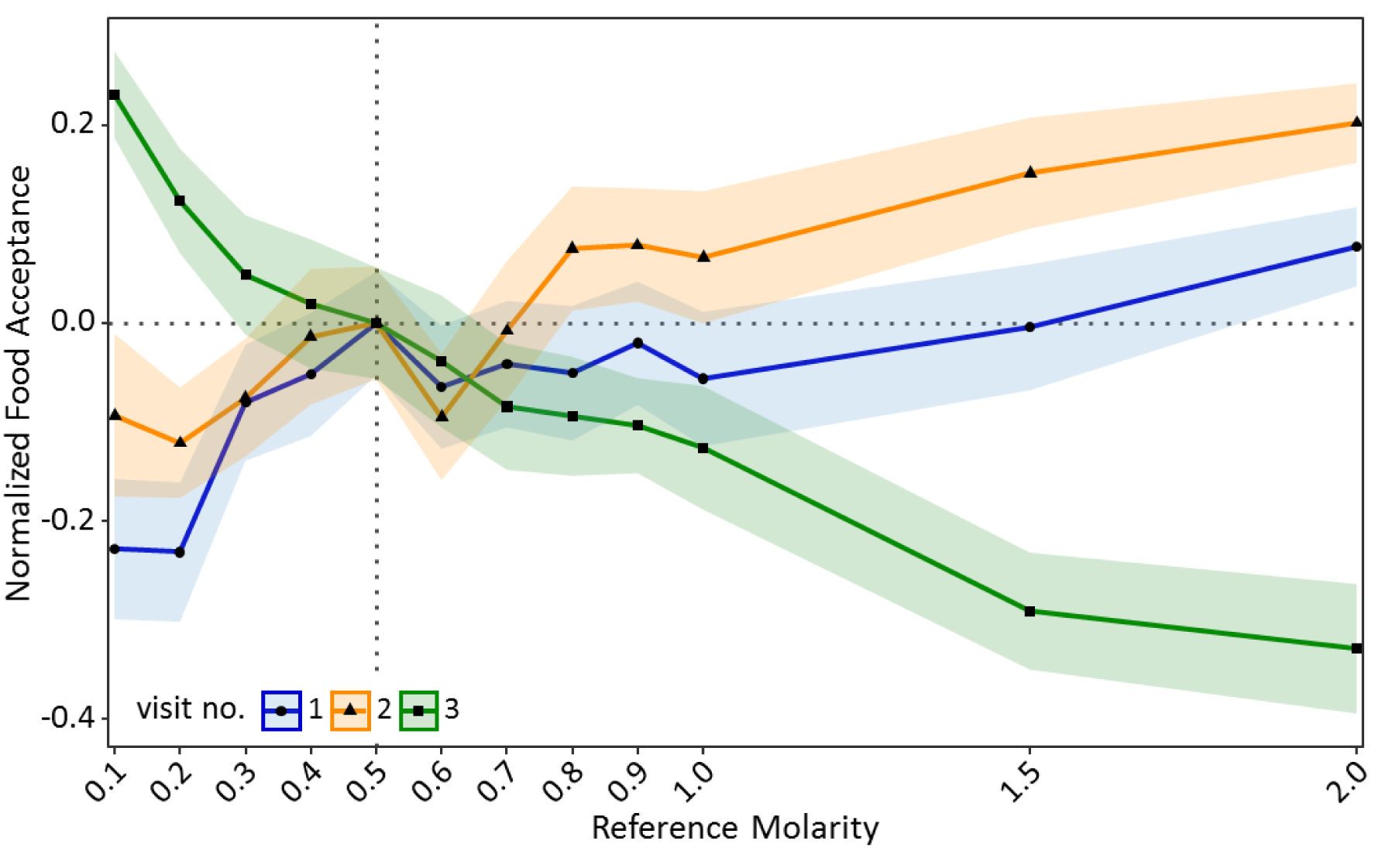
Food acceptance shown in experiment 1 for the two training visits (visit 1 & 2) in which ants received one of 12 molarities and the test visit (3) in which all ants received 0.5M. Shown are the mean food acceptance (points) and the 95% confidence intervals (coloured ribbons) for each reference molarity and visit. Data was normalized to show the mean food acceptance of the control group (received 0.5M on each visit) at 0 for all three visits. For a non-normalized graph of the data see supplement Figure S2.1.

A similar pattern was found for pheromone deposition behaviour on the way back to the nest (figure 3). In the training visits, number of pheromone depositions increased significantly with increasing molarity of the reference solution (GLMM: estimate= 0.86, z= 13.86, p<0.001). Additionally, ants performed significantly more pheromone depositions on the second return to the nest compared to the first return visit (GLMM: estimate= 0.31, z= 4.64, p<0.001). By contrast, on the test visit pheromone depositions decreased significantly with increasing molarity of the reference solution (GLMM: estimate= -0.82, z= -9.75, p<0.001, fig. 3). Ants which deposited more pheromone during the training visits generally deposited more pheromone on the test visit compared to ants which deposited less pheromone during the training visits (GLMM: estimate= 0.16, z= 15.99, p<0.001). Ants which were trained to very low molarities (0.2M: p<0.01) deposited significantly more pheromone depositions in the test visit than control ants, while ants trained to high molarities (1M: p<0.001, 1.5M: p<0.001, 2M: p<0.001) deposited less pheromone depositions than the control group (see supplement Table S2.2 for pairwise comparisons).

**Fig. 3:**
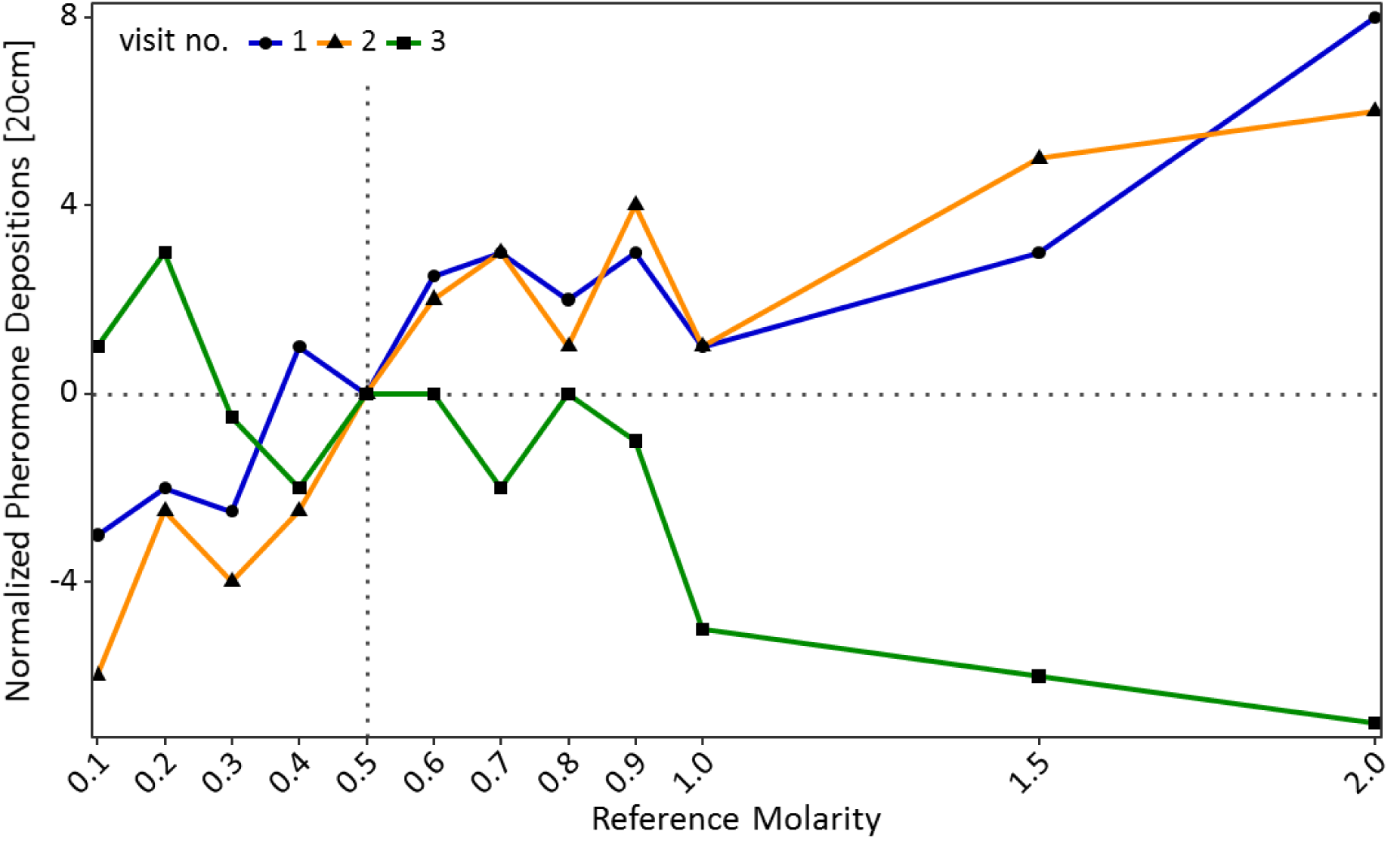
Pheromone depositions on the way back to the nest shown in experiment 1 for the two training visits (visit 1 & 2) in which ants received one of 12 molarities and the test visit (3) in which all ants received 0.5M. Shown is the median number of pheromone depositions (points) measured on a 20 cm track right behind the food source for each reference molarity and visit. Data was normalized to show the median number of pheromone depositions of the control group (received 0.5M on each visit) at 0 for all three visits. For a non-normalized graph of the data with error ribbons see supplement Figure S2.2.

## Experiment 2 – ruling out alternative explanations using scent training

The results of experiment 1 are consistent with relative value perception stemming from the psychological effects of successive contrasts. However, alternative hypotheses could also explain these results. Four possible alternatives must be excluded: sensory saturation, ingested sucrose changing haemolymph-sugar levels, psychophysical sensory contrast effects and the fact that ants may expect pre-shift solutions to return in later visits (see introduction). To rule out the above four alternative explanations, we carried out experiment 2.

### Experiment 2 - Methods

To rule out the alternative non-psychological explanations for the contrast effects we described above, we needed to change the expectation of the ants while exposing all ants to identical training regimes. This would provide a reference point for testing relative value perception while keeping sensory saturation, haemolymph-sugar levels, and psychophysical effects the same until the switch occurred. To this end, we trained ants over 8 visits to associate a high sucrose molarity (1.5M) with one scent, and a low molarity (0.25M) with a different scent. Then, in the 9^th^ testing phase, we used scents to trigger an expectation of either high or low molarity, which was then contrasted with a medium (0.5M) unscented solution. Finally, preference for the high-quality associated odour was tested for using a Y-maze.

For a detailed description of the methods used, see online supplement S3.1.

### Experiment 2 - Results

During training, ants behaved as expected, showing higher acceptance and pheromone deposition for 1.5M than 0.25M on all but the very first visit to 0.25M (Food acceptance: CLMM: estimate = -1.13, z= 3.38, p<0.001; pheromone depositions outward journey: GLMM: estimate= 1.79, z= 17.10, p<0.001; pheromone depositions inward journey: GLMM: estimate= -1.20, z= -10.10, p<0.001, figuress 4A, C & E). Furthermore, food acceptance and pheromone depositions both on the outward and inward journeys decreased with increasing experience with the 0.25M feeder and increased with increasing experience with the 1.5M feeder (Food acceptance: CLMM: estimate = -1.13, z= 3.38, p<0.001; pheromone depositions outward journey: GLMM: estimate= -0.31, z= -17.07, p<0.001; pheromone depositions inward journey: GLMM: estimate = -0.21, z= -7.02, p<0.001).

**Fig. 4:**
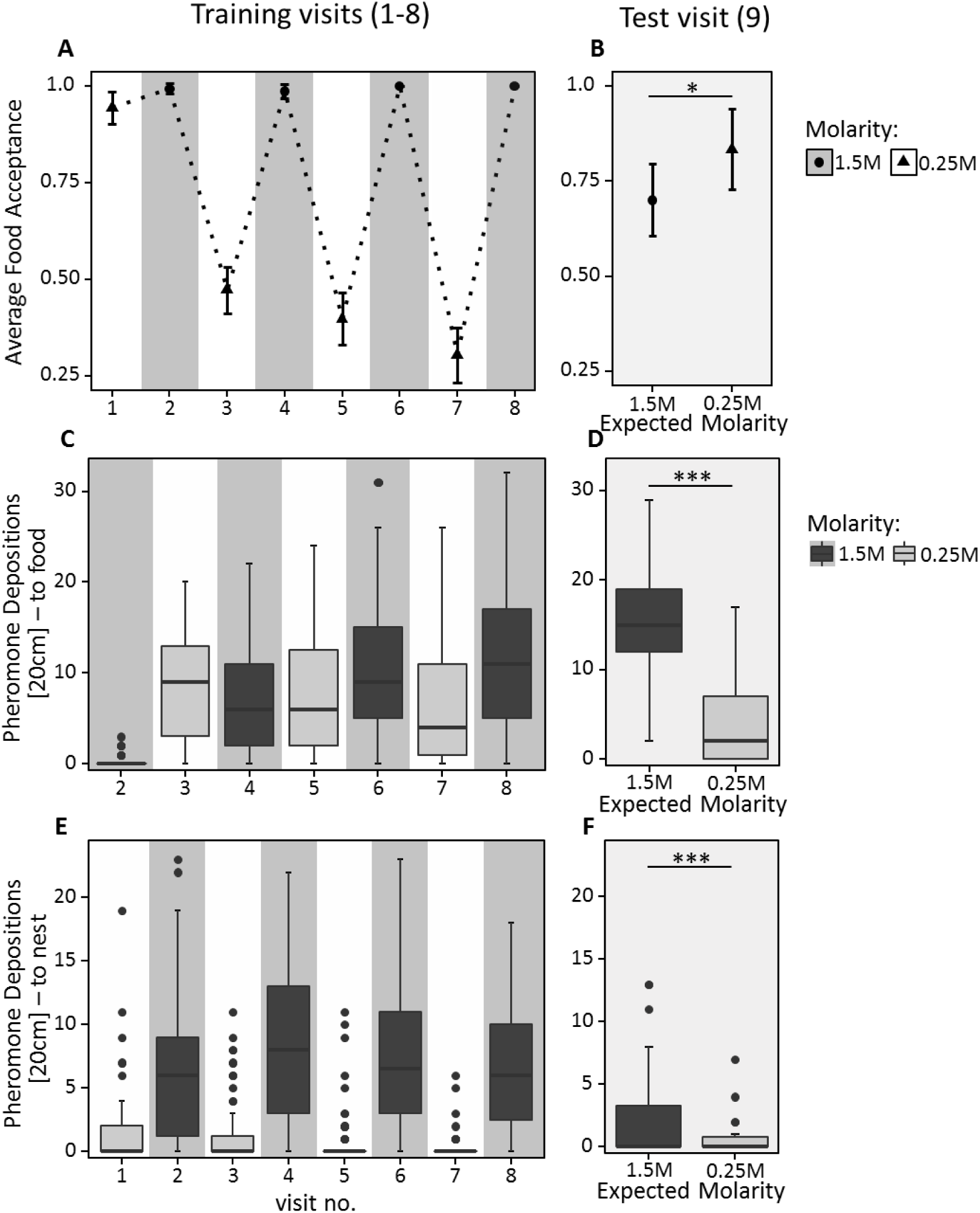
A) & B) Food acceptance C) & D) Number of pheromone depositions on the way to the food source and E) & F) Number of pheromone depositions on the way back to the nest shown in experiment 2 for A), C) & E) the eight training visits (visits 1-8) in which ants received 0.25M coupled with one scent and 1.5M coupled with another scent in an alternating order, always starting with 0.25M. and B), D) & F) the test visit (visit 9) in which ants always received unscented 0.5M sucrose solution, but the runway leading towards the food source was impregnated with one of the learned scents, triggering an expectation towards receiving either high or low molarities at the end of the runway. A) & B) Shown are the mean food acceptance (points) and the 95% confidence intervals (error bars) for each visit; C) - F) Shown are the median number of pheromone depositions on a 20 cm track right in front of the food source and the 75%/25% quantiles for each visit.

On the outward journey of the 9^th^ (test) visit, ants walking towards the feeder while exposed to 1.5M sucrose-associated cues deposited more pheromone (median=15, fig. 4D) compared to ants exposed to 0.25M-associated cues (median=2, GLMM: estimate= -1.32, z= -13.51, p<0.001). Moreover, in the learning probe, 87% of ants chose the 1.5M associated arm. This demonstrates that ants formed a robust expectation of food molarity based on the cues learned during training.

Ants exposed to 1.5M-associated cues during the 9^th^ visit showed significantly lower food acceptance towards the unscented 0.5M feeder than ants exposed to 0.25M-associated cues (CLMM: estimate= 1.07, z= 2.15, p= 0.03, figure 4B, table S1). Although ants exposed to high molarity associated cues on their outwards journey showed a significantly higher number of pheromone depositions on their return journey than ants confronted with low molarity scent (GLMM: estimate= -1.36, z= -5.50, p<0.001, figure 4E & F), the number of pheromone depositions decreased drastically for both treatments compared to training visits (median 1.5M = 0, median 0.25M = 0, figure 4E & F, table S1).

## Experiment 3 – expectation setting via trophallaxis: the nest as an information hub

Ants receive information about available food sources, such as food odour and palatability, through food exchanges (trophallaxis) inside the nest (Provecho and Josens 2009; Josens et al. 2016). An ant beginning a food scouting bout may not have direct information about the quality of the food sources available in the environment, but nonetheless must make a value judgement on their first visit to a food source. The aim of this experiment was to ascertain whether information about sucrose concentrations gained through trophallaxis in the nest affected the perceived value of food sources found outside the nest.

### Experiment 3 - Methods

An ant (forager) was allowed to feed at an unscented sucrose solution droplet of either 0.16, 0.5 or 1.5M at the end of a 60cm long runway. Once the ant had fed and returned to the nest, we observed the number of contacts with other nestmates which occurred until trophallaxis was initiated. When trophallaxis began, we noted the time spent in trophallaxis with the first trophallactic partner. When trophallaxis stopped, the receiving trophallactic partner (receiver) was gently moved from the nest and placed onto the start of a 20cm long runway, offering unscented 0.5M sucrose solution at the end. As the receiver fed, we noted its food acceptance.

### Experiment 3 - Results

The time spent in trophallaxis with the receiver increased significantly with increasing molarity (GLMM: estimate= 0.13, z= 4.79, p<0.001).

Acceptance scores of receivers towards 0.5M decreased with increasing molarity of the sucrose solution received through trophallaxis. The interaction of reference molarity and trophallaxis time significantly predicted acceptance (CLMM: estimate=-0.06, z= -2.34, p= 0.02, fig. 5). Ants which received 0.16M inside the nest showed significantly higher acceptance of 0.5M sucrose than ants which received 1.5M (p<0.01, see supplement Table S4.1 for pairwise comparisons).

**Fig. 5:**
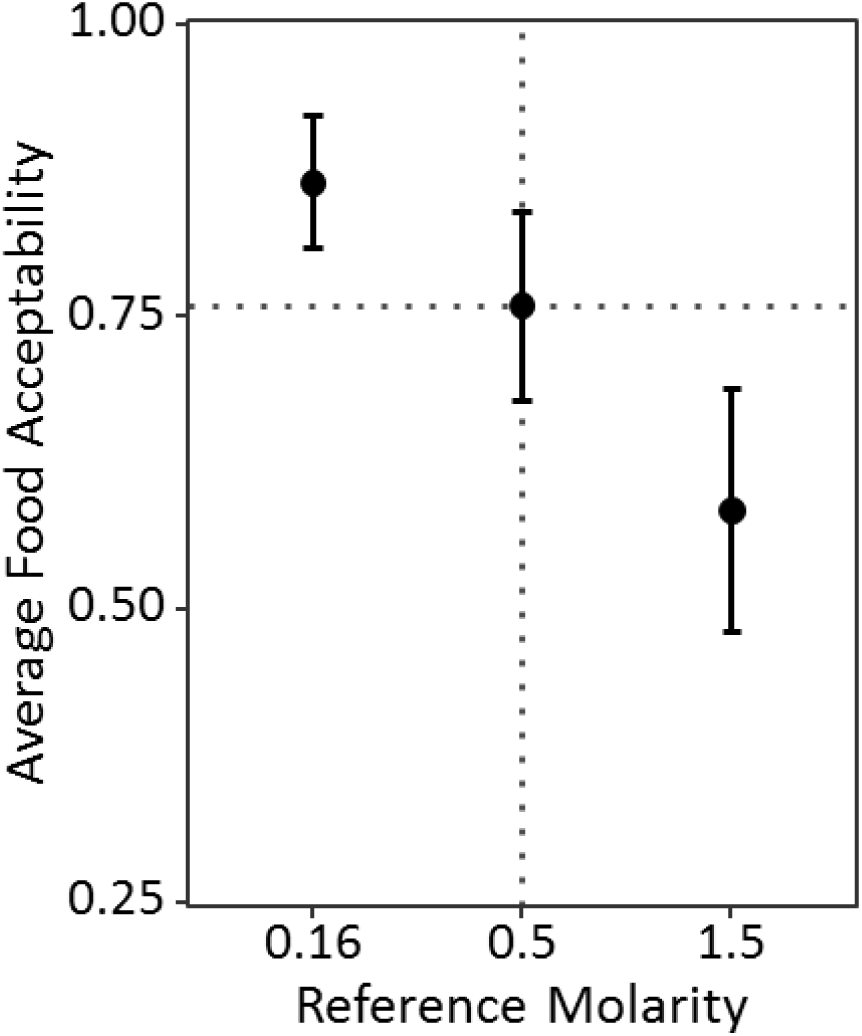
Food acceptance shown in experiment 3 for the receivers which received either 0.16, 0.5 or 1.5M through trophallaxis in the nest and then found 0.5M at the end of the runway. Shown are the mean food acceptance (points) and the 95% confidence intervals (error bars) for each reference molarity.

## Discussion

Kahneman and Tversky’s (1979) introduction of Prospect Theory contributed to a major shift in economic research by suggesting that humans do not perceive value in absolute terms, but relative to reference points. Here, we demonstrate parallel findings in an insect, providing for the first time to our knowledge a detailed description of relative value perception in an invertebrate. Positive contrast effects were shown by ants which were trained to low molarities (figures 2 & 3). These ants showed higher acceptance scores and deposited more pheromone after being shifted to medium quality than unshifted ants which received medium quality food throughout the whole experiment. Conversely, ants trained to high molarities showed lower acceptance after being shifted to medium quality compared to the unshifted control, showing negative contrast effects.

Another prediction of Prospect Theory, that gains are underemphasized and losses are overemphasized (Tversky and Kahneman 1992), is not supported by the data of our main experiment. Indeed, gains seem to be overvalued while losses are undervalued. This may be due to the psychophysics of our study system: a basic tenant of psychophysics is that the Just Noticeable Difference (JNDs) between two stimuli is a function of the relative difference between the stimuli (Fechner 1860; Stevens 1957; Zwislocki 2009). Thus, ants shifted from 0.1M to medium (0.5M) quality experience a 5-fold increase in molarity, while those down-shifted from 0.9M to 0.5M experience less than a two-fold decrease, although the absolute change was of the same magnitude. This would predict larger shift-changes, in terms of absolute molarity change, for gains than for losses. Indeed, the fact that this is also not seen may imply that losses are indeed – relatively speaking – looming larger than gains for the ants. Finally, it must be kept in mind that acceptance scores are unlikely to be linear, and that pheromone deposition behaviour shows large variation (Beckers, Deneubourg, and Goss 1992), making it difficult to use either of these factors to test for over- and undervaluation of gains and losses.

While the results of experiment 1 can be explained using alternative, non-psychological mechanisms (sensory saturation and changes in satiation) or rational behaviour based on future expectations, the results of experiment 2 cannot. Ants which were expecting high molarities after scent training showed lower acceptance scores when confronted with unscented medium quality food than ants which expected to find low quality food (figure 4B). This is in spite of all ants undergoing identical training experiences. The only difference between the groups was the odour of the runway on the 9^th^ (test) visit. It is thus unlikely that sensory saturation, increased haemolymph-sugar levels, simple psychophysical effects or expecting pre-shift solutions to return can fully explain the behaviour of the ants in our experiments.

Contrast effects were stronger in experiment 1 than in experiment 2. Possible explanations for this pattern are given in supplementary note 1. The fact that we nonetheless see both positive and negative contrasts suggests that such contrast effects are very pronounced. The reduced pheromone deposition seen in the final return in experiment 2 may be due to the change in environment (scented runways to unscented runways) causing a disruption in recruitment behaviour, perhaps due to generalization decrement (E. D. Capaldi 1978; Kimble 1961) or neophobia (Barnett 1958; Johnson 2000, 2000; Mitchell 1976; Pliner and Loewen 1997).

Ants which received information about the quality of a food source through trophallactic interactions inside the nest are able to use this information when evaluating new food sources outside the nest. Ants which received low quality (0.16M) food from a returning forager were more likely to accept medium (0.5M) food when foraging themselves than ants which had received good (1.5M) food via trophallaxis (fig. 5). Apart from ants valuing the medium quality food source based only on the quality they received from the returning forager, there is another possible explanation which may lead to the same pattern of food acceptance as shown in this experiment (fig. 5). Ants which expected to find a high quality food source outside the nest may not have accepted a medium quality food source in order to search for the high quality food source which is supposed to be available outside the nest, leading to low food acceptance scores when the reference point was high (Wendt and Czaczkes 2017). Our results suggest that information about sucrose concentrations gained through trophallactic interactions inside the nest can affect the way a newly discovered food source is valued outside the nest. Trophallaxis is a rich source of information: it has been shown to contain chemical cues, growth proteins, and hormones (LeBoeuf et al. 2016). Transfer of scented food (Provecho and Josens 2009; Josens et al. 2016) and aphid-associated information (Hayashi et al. 2017) through trophallactic contacts inside the nest, as well as information about available food qualities gained directly or through pheromone trails (Beckers, Lachaud, and Fresneau 1994; Czaczkes and Beckwith 2018; Roces and Núñez 1993; Roces 1993; Wendt and Czaczkes 2017) have been shown to shape ant behaviour outside the nest. By taking into account information gained inside the nest, recruited workers will be able to evaluate newly discovered food sources in relation to other food sources available in the environment. They will also be able to make better informed decisions on whether it is worth exploiting a new food source or ignore it. Such a pattern would lead to individual ants being more likely to forego food sources which are of lower quality than the average available food sources and thus allows colonies to only exploit above-average food sources. Ants can also use this information to choose between various information use strategies, such as whether to continue exploiting known food sources or be recruited to follow pheromone trails leading to other food sources (Czaczkes and Beckwith 2018). Ultimately, we see the nest serving as an information hub, in which information about currently available food sources can be collected, synthesised, and fed back to outgoing foragers. Relative value perception can therefore be expected to have strong effects not only on the individual behaviour of animals, but also on the collective behaviour of insect colonies, potentially allowing colonies to ignore usually acceptable options in favour of better ones

A broad range of behaviours relevant to behavioural economics have now been described in invertebrates. These include overvaluing rewards in which more effort was invested (Czaczkes et al. 2018), self-control (Cheng et al. 2002; Wendt and Czaczkes 2017), and state dependent learning (Pompilio, Kacelnik, and Behmer 2006). Many other parallels to human behaviour and cognition have also been described in insects, such as abstract association learning (Czaczkes et al. 2014; Giurfa, Eichmann, and Menzel 1996; Hateren, Srinivasan, and Wait 1990), concept learning (Giurfa et al. 2001), and reward changes affecting voluntary task switching (Czaczkes et al. 2018). Applying concepts from behavioural economics to the study of animal behaviour is likely to yield many further insights. Moreover, the benefits of an interdisciplinary approach are likely to flow both ways. We suggest that invertebrates make attractive models for a broader understanding of behavioural economics in humans. Using animal models allows researchers to avoid pitfalls associated with studies on humans, such as cultural and educational differences (Carter and Irons 1991; Guiso, Sapienza, and Zingales 2006) second-guessing of experimenters, and non-relevant reward sizes (Levitt and List 2007) as well as relaxing ethical concerns.

Due to its complexity, building models which can accurately predict human behaviour is a challenge. This is compounded by the fact that data on humans obtained in laboratory experiments overwhelmingly stem from game-like designs that are highly artificial and where the economic incentives that can be provided to experimental subjects are severely limited by the research budget of the experimenter (Kahneman and Tversky 1979; Levitt and List 2007). At the same time, there has been much progress in field studies on humans to clearly measure causal relationships (Harrison and List 2004). However, the usefulness of these new techniques (such as field experiments) is clearly constrained by the range of questions and settings to which they can be meaningfully applied. Hence, while behavioural studies on invertebrates also have their limitations (for example, in that inducing expectations is more of a challenge), they can be easily designed to be ecologically meaningful, and offer rewards which are in line with the real-life budgets under which the animals operate. Therefore, we propose that economic models to predict invertebrate decision making may be a complementary step on the way to predict human behaviour.

While there is a well-developed tradition of integrating economics and biology (Aw et al. 2009; Aw, Vasconcelos, and Kacelnik 2011; Cheng et al. 2002; Czaczkes et al. 2018; Evans and Westergaard 2006; Lydall, Gilmour, and Dwyer 2010; Wendt and Czaczkes 2017), we feel a critical mass of evidence is now available to consider comparative behavioural economics as a relevant discipline for both biologists and economists.

## Acknowledgements

We thank Flavio Roces for helpful comments on this work and Florian Hartig for advice concerning statistical analysis of our data.

## References

Annicchiarico, Ivan, Amanda C. Glueck, Lucas Cuenya, Katsuyoshi Kawasaki, Shannon E. Conrad, and Mauricio R. Papini. 2016. “Complex Effects of Reward Upshift on Consummatory Behavior.” Behavioural Processes 129 (August): 54–67. https://doi.org/10.1016/j.beproc.2016.06.006.

Aw, J. M., R. I. Holbrook, T. Burt de Perera, and A. Kacelnik. 2009. “State-Dependent Valuation Learning in Fish: Banded Tetras Prefer Stimuli Associated with Greater Past Deprivation.” Behavioural Processes, Proceedings of the meeting of the Society for the Quantitative Analyses of Behavior (SQAB 2008), which took place in Chicago IL, May 22-24 of 2008, 81 (2): 333–36. https://doi.org/10.1016/j.beproc.2008.09.002.

Aw, J. M., Marco Vasconcelos, and Alex Kacelnik. 2011. “How Costs Affect Preferences: Experiments on State Dependence, Hedonic State and within-Trial Contrast in Starlings.” Animal Behaviour 81 (6): 1117–28. https://doi.org/10.1016/j.anbehav.2011.02.015.

Barnett, S. A. 1958. “Experiments on ‘Neophobia’ in Wild and Laboratory Rats.” British Journal of Psychology 49 (3): 195–201. https://doi.org/10.1111/j.2044-8295.1958.tb00657.x.

Beckers, R., J. L. Deneubourg, and S. Goss. 1992. “Trail Laying Behaviour during Food Recruitment in the Ant Lasius Niger (L.).” Insectes Sociaux 39: 59–72.

Beckers, R., J. L. Deneubourg, and S. Goss. 1993. “Modulation of Trail Laying in the AntLasius Niger (Hymenoptera: Formicidae) and Its Role in the Collective Selection of a Food Source.” Journal of Insect Behavior 6 (6): 751–59. https://doi.org/10.1007/BF01201674.

Beckers, R., J. P. Lachaud, and D. Fresneau. 1994. “The Influence of Olfactory Conditioning on Food Preference in the Ant Lasius Niger (L.).” Ethology Ecology & Evolution 6 (2): 159–67. https://doi.org/10.1080/08927014.1994.9522991.

Bentosela, Mariana, Adriana Jakovcevic, Angel M. Elgier, Alba E. Mustaca, and Mauricio R. Papini. 2009. “Incentive Contrast in Domestic Dogs (Canis Familiaris).” Journal of Comparative Psychology 123 (2): 125–30. https://doi.org/10.1037/a0013340.

Bitterman, M.E. 1976. “Incentive Contrast in Honey Bees.” Science 192 (4237): 380–82.

Black, R. W. 1968. “Shifts in Magnitude of Reward and Contrast Effects in Instrumental and Selective Learning: A Reinterpretation.” Psychological Review 75 (2): 114–26. https://doi.org/10.1037/h0025563.

Bower, G. H. 1961. “A Contrast Effect in Differential Conditioning.” Journal of Experimental Psychology 62 (2): 196–99. http://dx.doi.org/10.1037/h0048109.

Boyce, Christopher J., Gordon D. A. Brown, and Simon C. Moore. 2010. “Money and Happiness: Rank of Income, Not Income, Affects Life Satisfaction.” Psychological Science 21 (4): 471–75. https://doi.org/10.1177/0956797610362671.

Brosnan, Sarah F., and Frans B. M. de Waal. 2003. “Monkeys Reject Unequal Pay.” Nature 425 (6955): 297. https://doi.org/10.1038/nature01963.

Campbell, Patrick E., Charles M. Crumbaugh, Stephen B. Knouse, and M. Emily Snodgrass. 1970. “A Test of the ‘;Ceiling Effect’; Hypothesis of Positive Contrast.” Psychonomic Science 20 (1): 17–18. https://doi.org/10.3758/BF03335577.

Capaldi, E. J., and D. Lynch. 1967. “Repeated Shifts in Reward Magnitude: Evidence in Favor of an Associational and Absolute (Noncontextual) Interpretation.” Journal of Experimental Psychology 75 (2): 226–35. https://doi.org/10.1037/h0024986.

Capaldi, Elizabeth D. 1978. “Effects of Changing Alley Color on the Successive Negative Contrast Effect.” Bulletin of the Psychonomic Society 12 (1): 69–70. https://doi.org/10.3758/BF03329628.

Carter, John R., and Michael D. Irons. 1991. “Are Economists Different, and If So, Why?” Journal of Economic Perspectives 5 (2): 171–77. https://doi.org/10.1257/jep.5.2.171.

Cheng, Ken, Jennifer Peña, Melanie A. Porter, and Julia D. Irwin. 2002. “Self-Control in Honeybees.” Psychonomic Bulletin & Review 9 (2): 259–63. https://doi.org/10.3758/BF03196280.

Couvillon, P.A., and M.E. Bitterman. 1984. “The Overlearning-Extinction Effect and Succeessive Contrasts in Honeybees.” Journal of Comparative Psychology 98 (1): 100–109.

Crespi, Leo P. 1942. “Quantitative Variation of Incentive and Performance in the White Rat.” The American Journal of Psychology55 (4): 467–517. https://doi.org/10.2307/1417120.

Czaczkes, Tomer J., and John J. Beckwith. 2018. “Information Synergy: Adding Unambiguous Quality Information Rescues Social Information Use in Ants.” BioRxiv, February, 219980. https://doi.org/10.1101/219980.

Czaczkes, Tomer J., Birgit Brandstetter, Isabella di Stefano, and Jürgen Heinze. 2018. “Greater Effort Increases Perceived Value in an Invertebrate.” Journal of Comparative Psychology, March.

Czaczkes, Tomer J., Christoph Grüter, Laura Ellis, Elizabeth Wood, and Francis L. W. Ratnieks. 2013. “Ant Foraging on Complex Trails: Route Learning and the Role of Trail Pheromones in Lasius Niger.” Journal of Experimental Biology 216 (2): 188–97. https://doi.org/10.1242/jeb.076570.

Czaczkes, Tomer J., Christoph Grüter, and Francis L. W. Ratnieks. 2015. “Trail Pheromones: An Integrative View of Their Role in Social Insect Colony Organization.” Annual Review of Entomology 60 (1): 581–99. https://doi.org/10.1146/annurev-ento-010814-020627.

Czaczkes, Tomer J., Alexandra Koch, K. Fröber, and G. Dreisbach. 2018. “Voluntary Switching in an Invertebrate: The Effect of Cue and Reward Change.” Journal of Experimental Psychology: Animal Learning and Cognition.

Czaczkes, Tomer J., Linda Schlosser, Jürgen Heinze, and Volker Witte. 2014. “Ants Use Directionless Odour Cues to Recall Odour-Associated Locations.” Behavioral Ecology and Sociobiology 68 (6): 981–88. https://doi.org/10.1007/s00265-014-1710-2.

Devigne, C., and C. Detrain. 2002. “Collective Exploration and Area Marking in the Ant <Emphasis Type=“Italic”>Lasius Niger</Emphasis>.” Insectes Sociaux 49 (4): 357–62. https://doi.org/10.1007/PL00012659.

Dunham, P. J. 1968. “Contrasted Conditions of Reinforcement: A Selective Critique.” Psychological Bulletin 69 (5): 295–315. https://doi.org/10.1037/h0025690.

Dussutour, Audrey, Vincent Fourcassié, Dirk Helbing, and Jean-Louis Deneubourg. 2004. “Optimal Traffic Organization in Ants under Crowded Conditions.” Nature 428 (6978): 70–73. https://doi.org/10.1038/nature02345.

Evans, T.A., and G.C. Westergaard. 2006. “Self Control and Tool Use in Tufted Capuchin Monkeys.” Journal of Comparative Psychology 120 (2): 163–66. https://doi.org/10.1037/0735-7036.120.2.163.

Fechner, G. T. 1860. Elemente Der Psychophysik [Elements of Psychophysics]. Vol. Band 2. Leipzig: Breitkopf und Härtel.

Flaherty, Charles F. 1982. “Incentive Contrast: A Review of Behavioral Changes Following Shifts in Reward.” Animal Learning & Behavior 10 (4): 409–40. https://doi.org/10.3758/BF03212282.

Flaherty, Charles F.. 1999. Incentive Relativity. Problems in the Behavioural Sciences 15. Cambridge University Press.

Flaherty, Charles F., Howard C. Becker, and Larissa Pohorecky. 1985. “Correlation of Corticosterone Elevation and Negative Contrast Varies as a Function of Postshift Day.” Animal Learning & Behavior 13 (3): 309–14. https://doi.org/10.3758/BF03200025.

Giurfa, Martin, Birgit Eichmann, and Randolf Menzel. 1996. “Symmetry Perception in an Insect.” Nature 382 (6590): 458–61. https://doi.org/10.1038/382458a0.

Giurfa, Martin, Shaowu Zhang, Arnim Jenett, Randolf Menzel, and Mandyam V. Srinivasan. 2001. “The Concepts of ‘Sameness’ and ‘Difference’ in an Insect.” Nature 410 (6831): 930–33. https://doi.org/10.1038/35073582.

Guiso, Luigi, Paola Sapienza, and Luigi Zingales. 2006. “Does Culture Affect Economic Outcomes?” Journal of Economic Perspectives 20 (2): 23–48. https://doi.org/10.1257/jep.20.2.23.

Hangartner, W. 1970. “Control of Pheromone Quantity in Odor Trails of the Ant Acanthomyops Interjectus.” Experientia 26 (6): 664–65. https://doi.org/10.1007/BF01898753.

Harrison, Glenn W., and John A. List. 2004. “Field Experiments.” Journal of Economic Literature 42 (4): 1009–55. https://doi.org/10.1257/0022051043004577.

Hateren, J. H. van, M. V. Srinivasan, and P. B. Wait. 1990. “Pattern Recognition in Bees: Orientation Discrimination.” Journal of Comparative Physiology A 167 (5): 649–54. https://doi.org/10.1007/BF00192658.

Hayashi, Masayuki, Masaru K. Hojo, Masashi Nomura, and Kazuki Tsuji. 2017. “Social Transmission of Information about a Mutualist via Trophallaxis in Ant Colonies.” Proc. R. Soc. B 284 (1861): 20171367. https://doi.org/10.1098/rspb.2017.1367.

Helson, H. 1964. Adaptation-Level Theory: An Experimental and Systematic Approach to Behavior. New York: Harper and Row.

Holman, Luke, Megan L. Head, Robert Lanfear, and Michael D. Jennions. 2015. “Evidence of Experimental Bias in the Life Sciences: Why We Need Blind Data Recording.” PLOS Biology. https://doi.org/10.1371/journal.pbio.1002190.

Johnson, Elizabeth. 2000. “Food-Neophobia in Semi-Free Ranging Rhesus Macaques: Effects of Food Limitation and Food Source.” American Journal of Primatology 50 (1): 25–35. https://doi.org/10.1002/(SICI)1098-2345(200001)50:1<25::AID-AJP3>3.0.CO;2-D.

Josens, Roxana, Analia Mattiacci, Jimena Lois-Milevicich, and Alina Giacometti. 2016. “Food Information Acquired Socially Overrides Individual Food Assessment in Ants.” Behavioral Ecology and Sociobiology 70 (12): 2127–38. https://doi.org/10.1007/s00265-016-2216-x.

Josens, Roxana, and Flavio Roces. 2000. “Foraging in the Ant Camponotus Mus: Nectar-Intake Rate and Crop Filling Depend on Colony Starvation.” Journal of Insect Physiology 46 (7): 1103–10. https://doi.org/10.1016/S0022-1910(99)00220-6.

Kahneman, Daniel, and Amos Tversky. 1979. “Prospect Theory: An Analysis of Decision under Risk.” Econometrica 47 (2): 263–91.

Kimble, G. A. 1961. Hilgard and Marquis’ “Conditioning and Learning.” Vol. 2nd edition. East Norwalk, CT, US: Appleton-Century-Crofts.

LeBoeuf, Adria C, Patrice Waridel, Colin S Brent, Andre N Gonçalves, Laure Menin, Daniel Ortiz, Oksana Riba-Grognuz, et al. 2016. “Oral Transfer of Chemical Cues, Growth Proteins and Hormones in Social Insects.” ELife 5. https://doi.org/10.7554/eLife.20375.

Levitt, Steven D., and John A. List. 2007. “What Do Laboratory Experiments Measuring Social Preferences Reveal About the Real World?” Journal of Economic Perspectives 21 (2): 153–74. https://doi.org/10.1257/jep.21.2.153.

Lydall, Emma S., Gary Gilmour, and Dominic M. Dwyer. 2010. “Rats Place Greater Value on Rewards Produced by High Effort: An Animal Analogue of the ‘Effort Justification’ Effect.” Journal of Experimental Social Psychology 46 (6): 1134–37. https://doi.org/10.1016/j.jesp.2010.05.011.

Mailleux, A.-C., C. Detrain, and J. L. Deneubourg. 2006. “Starvation Drives a Threshold Triggering Communication.” The Journal of Experimental Biology 209: 4224–29. https://doi.org/10.1242/jeb.02461.

Mankiw, N. G. 2011. Principles of Economics. South-Western.

Mayer-Gross, W., and J. W. Walker. 1946. “Taste and Selection of Food in Hypoglycaemia.” British Journal of Experimental Pathology 27 (5): 297–305.

McBride, R. L. 1982. “Range Bias in Sensory Evaluation.” International Journal of Food Science & Technology 17 (3): 405–10. https://doi.org/10.1111/j.1365-2621.1982.tb00195.x.

McNamara, John M., Tim W. Fawcett, and Alasdair I. Houston. 2013. “An Adaptive Response to Uncertainty Generates Positive and Negative Contrast Effects.” Science 340 (6136): 1084–86. https://doi.org/10.1126/science.1230599.

Melanson, Kathleen J., Margriet S. Westerterp-Plantenga, L. Arthur Campfield, and Wim H. M. Saris. 1999. “Blood Glucose and Meal Patterns in Time-Blinded Males, after Aspartame, Carbohydrate, and Fat Consumption, in Relation to Sweetness Perception.” British Journal of Nutrition 82 (6): 437–46. https://doi.org/10.1017/S0007114599001695.

Mitchell, D. 1976. “Experiments on Neophobia in Wild and Laboratory Rats: A Reevaluation.” Journal of Comparative and Physiological Psychology 90 (2): 190–97. http://dx.doi.org/10.1037/h0077196.

Mustaca, Alba E., Mariana Bentosela, and Mauricio R. Papini. 2000. “Consummatory Successive Negative Contrast in Mice.” Learning and Motivation 31 (3): 272–82. https://doi.org/10.1006/lmot.2000.1055.

Neumann, J. von, and O. Morgenstern. 1944. Theory of Games and Economic Behavior. Princeton, NJ: Princeton University Press.

Núñez, Josué A. 1966. “Quantitative Beziehungen zwischen den Eigenschaften von Futterquellen und dem Verhalten von Sammelbienen.” Zeitschrift für vergleichende Physiologie 53 (2): 142–64. https://doi.org/10.1007/BF00343733.

Papini, Mauricio R., H. Wayne Ludvigson, David Huneycutt, and Robert L. Boughner. 2001. “Apparent Incentive Contrast Effects in Autoshaping with Rats.” Learning and Motivation 32 (4): 434–56. https://doi.org/10.1006/lmot.2001.1088.

Parducci, Allen. 1984. “Value Judgments: Toward a Relational Theory of Happiness.” In Attitudinal Judgment, 3–21. Springer Series in Social Psychology. Springer, New York, NY. https://doi.org/10.1007/978-1-4613-8251-5_1.

Pellegrini, Santiago, and Alba Mustaca. 2000. “Consummatory Successive Negative Contrast with Solid Food.” Learning and Motivation 31 (2): 200–209. https://doi.org/10.1006/lmot.2000.1052.

Pliner, PATRICIA, and E. RUTH Loewen. 1997. “Temperament and Food Neophobia in Children and Their Mothers.” Appetite 28 (3): 239–54. https://doi.org/10.1006/appe.1996.0078.

Pompilio, Lorena, Alex Kacelnik, and Spencer T. Behmer. 2006. “State-Dependent Learned Valuation Drives Choice in an Invertebrate.” Science 311 (5767): 1613–15. https://doi.org/10.1126/science.1123924.

Premack, D., and W. A. Hillix. 1962. “Evidence for Shift Effects in the Consummatory Response.” Journal of Experimental Psychology 63 (3): 284–88. https://doi.org/10.1037/h0039368.

Provecho, Yael, and Roxana Josens. 2009. “Olfactory Memory Established during Trophallaxis Affects Food Search Behaviour in Ants.” Journal of Experimental Biology 212 (20): 3221–27. https://doi.org/10.1242/jeb.033506.

Riskey, Dwight R., Allen Parducci, and Gary K. Beauchamp. 1979. “Effects of Context in Judgments of Sweetness and Pleasantness.” Perception & Psychophysics 26 (3): 171–76. https://doi.org/10.3758/BF03199865.

Roces, Flavio. 1993. “Both Evaluation of Resource Quality and Speed of Recruited Leaf-Cutting Ants (Acromyrmex Lundi) Depend on Their Motivational State.” Behavioral Ecology and Sociobiology 33 (3): 183–89. https://doi.org/10.1007/BF00216599.

Roces, Flavio, and J. A. Núñez. 1993. “Information about Food Quality Influences Load-Size Selection in Recruited Leaf-Cutting Ants.” Animal Behaviour 45 (1): 135–43. https://doi.org/10.1006/anbe.1993.1012.

Stevens, S.S. 1957. “On the Psychophysical Law.” The Psychological Review 64 (3): 153–81.

Tinklepaugh, L. O. 1928. “An Experimental Study of Representative Factors in Monkeys.” Journal of Comparative Psychology 8 (3): 197–236. https://doi.org/10.1037/h0075798.

Tversky, Amos, and Daniel Kahneman. 1974. “Judgment under Uncertainty: Heuristics and Biases.” Science 185 (4157): 1124–31. https://doi.org/DOI:10.1126/science.185.4157.1124.

Tversky, Amos, and Daniel Kahneman. 1981. “The Framing of Decisions and the Psychology of Choice.” Science 211 (4481): 453–58. https://doi.org/10.1126/science.7455683.

Tversky, Amos, and Daniel Kahneman. 1992. “Advances in Prospect Theory: Cumulative Representation of Uncertainty.” Journal of Risk and Uncertainty 5 (4): 297–323. https://doi.org/10.1007/BF00122574.

Ungemach, Christoph, Neil Stewart, and Stian Reimers. 2011. “How Incidental Values From the Environment Affect Decisions About Money, Risk, and Delay.” Psychological Science 22 (2): 253–60. https://doi.org/10.1177/0956797610396225.

Vlaev, Ivo, Nick Chater, Neil Stewart, and Gordon D. A.Brown. 2011. “Does the Brain Calculate Value?” Trends in Cognitive Sciences 15 (11): 546–54. https://doi.org/10.1016/j.tics.2011.09.008.

Vogel, J. R., P. J. Mikulka, and N. E. Spear. 1968. “Effects of Shifts in Sucrose and Saccharine Concentrations on Licking Behavior in the Rat.” Journal of Comparative and Physiological Psychology 66 (3): 661–66. https://doi.org/10.1037/h0026556.

Weinstein, Lawrence. 1970a. “Negative Incentive Contrast with Sucrose.” Psychonomic Science 19 (1): 13–14. https://doi.org/10.3758/BF03335483.

Weinstein, Lawrence. 1970b. “Negative Incentive Contrast Effects with Saccharin vs Sucrose and Partial Reinforcement.” Psychonomic Science 21 (5): 276–78. https://doi.org/10.3758/BF03330713.

Wendt, Stephanie, and Tomer J. Czaczkes. 2017. “Individual Ant Workers Show Self-Control.” Biology Letters 13 (10): 20170450. https://doi.org/10.1098/rsbl.2017.0450.

Zwislocki, Jozef J. 2009. Sensory Neuroscience: Four Laws of Psychophysics. Springer US.

